# Modulation of macrophage defense responses by Mycobacterial persistence protein MprA (*Rv0981*) in human THP-1 cells: effect of single amino acid variation on host-pathogen interactions

**DOI:** 10.1101/2020.04.27.063602

**Authors:** Kausik Bhattacharyya, Upasana Bandopadhyay, Aayushi Singh, Amresh Prakash, Vishal Nemaysh, Shruti Jain, Mandira Varma-Basil, Andrew M Lynn, Mridula Bose, Pratibha Mehta Luthra, Krishnamurthy Natarajan, Vani Brahmachari

## Abstract

*M. tuberculosis* is one of the most successful human pathogens causing tuberculosis that leads to highest daily morbidity worldwide. The evasion of the host immune responses is an important strategy that *M. tuberculosis* adopts. MprA (*Rv0981*), the response regulator of two component system is known for DNA binding activity in the pathogen and its role in persistent infection in the host. MprA is recognized as a late stage antigen during infection. A variant form of the protein MprA with G70S polymorphism (MprA*) is observed in one of our local and in several global clinical isolates of *M. tuberculosis.* Here we report the nuclear localization of MprA and MprA* in differentiated macrophages. MprA and MprA* increase the expression of TGF-β and IL-10, the immune suppressive cytokines in THP-1 derived macrophage cells. Concurrently the phago-lysosome fusion is significantly reduced as shown by infection with *M.bovis* BCG. We show that single nucleotide variation in clinical isolates lead to quantitative variations resulting in host immune suppression and support the survival and persistence of the pathogen.

## Introduction

Tuberculosis is one of the deadliest communicable diseases that infects one-third of the population. *M.tuberculosis* has adopted several immune escape mechanisms that prevent the host from clearing the pathogen. A plethora of immune cells migrate to the site of infection in lungs during early stages of infection (Marshall et al., 2018).

MprA(*Rv0981*) is one of the proteins known to regulate more than 200 genes in *M.tuberculosis* (Pang et al., 2007; Bretl et al., 2011). It is identified as an important hub in protein interactome network of *M. tuberculosis* (Vashisht et al., 2012) and is an essential gene in *M tuberculosis* as demonstrated by transposon mutagenesis (Sassetti et al., 2003). MprA is a response regulator and with its counterpart MprB (*Rv0982*), the sensor kinase, functions as a two-component signal transduction system. It is involved in the regulation of various stress-responsive genes, including the up-regulation of two sigma factors *sigE(Rv1221)* and sigB(*Rv2710*)(He et al., 2006). It is also responsible for the repression of multiple genes of the hypoxia associated regulons, iron metabolism and starvation and is required for the establishment and persistence of infection inside the host cells (Zahrt & Deretic, 2001; He et al., 2006). Interestingly, MprA is known to be expressed at day 5 of infection in THP-1 macrophages and is detected in the lungs of tuberculosis infected mice at 15 days post-infection (Gupta et al., 2010).Several studies on *Mycobacterium* clinical isolates have shown the occurrence of sequence variations in many important gene loci and in some cases these variations are known to affect pathogenesis, drug resistance, virulence and immunity (Cubillos-Ruiz et al., 2008; Zakham et al., 2012).

In one of the multi-drug resistant (MDR) Indian clinical isolate VPCI591, we found a single nucleotide variation in the *mprA* gene (G70S; NCBI: SRX5802345, BioSample:SAMN11568242) and the same non-synonymous variation was detected in the global data with considerable frequency (3%). In the light of the antigenic potential of MprA, we examined the cellular localization and the effect of the SNV in THP-1 cells. We found that non-synonymous variation (G70S) alters the host-immune suppression and brings about structural variation in MprA protein.

## Experimental procedures

### In-silico analysis

The sequence of clinical isolate VPCI591(PRJNA540936), from India was analysed using Maq(Li et al., 2008), inGAP(Qi et al., 2009) and GATK pipeline(McKenna et al., 2010) and G70S variation in MprA was detected and designated as MprA *(Bhattacharyya et al., 2020). The global frequency of MprA* was calculated from GMTV database (http://mtb.dobzhanskycenter.org) (Chernyaeva et al., 2014) and tbVar database (http://genome.igib.res.in/tbvar/) (Joshi et al., 2014).

### Cloning and Expression of MprA and MprA*(G70S)

*M. tuberculosis* H37Rv was maintained on “Lowenstein–Jensen” (LJ) medium and were grown in Middlebrook 7H9 broth (Becton Dickinson) supplemented with 10 % OADC (Himedia FD329) filtered 0.2% glycerol (SRL) and 0.05% Tween 80 (Sigma) at 37°C. The genomic DNA from *M. tuberculosis* H37Rv was isolated using CTAB (cetyltrimethyl ammonium bromide) method (Bose et al., 1993). The concentration and purity of the isolated DNA was measured using Qubit spectrophotometer (Invitrogen) and further quality was assessed by electrophoresis on 0.8% agarose gel. MprA gene was amplified by PCR using Phusion high fidelity polymerase (Fermentas) with the specific primers (Rv0981_Fp-GTGCGAATTCTTGTCG, Rv0981_Rp-TCAGGGTGGTGTTTC)QIAquick Gel Extraction kit (Qiagen) was used to extract the amplicon and was cloned in pET28a+ vector (Novagen). We carried out site-directed mutagenesis to insert the G70S variation with suitable primers using Agilent site directed mutagenesis kit (Quickchange II).The insert sequence in the clones confirmed by sequencing. The histidine-tagged proteins MprA and MprA* were purified using Ni-NTA Agarose beads (Qiagen) using standardized protocol(Chadha et al., 2015; Mehto et al., 2015). The protein was isolated from inclusion bodies from the culture of *E.coli* BL21(DE3) induced for expression of MprA and MprA*, dissolved in 8M Urea and dialyzed against 1M Urea. The protein fraction was loaded on the beads and the flow through after washing with 10mM imidazole (wash1) and 20mM imidazole (wash 2) were collected. The 6Xhis-MprA/MprA* was eluted with 250mM imidazole and analysed by 15% PAGE (Supplementery Fig 1).

### Cell Culture and protein stimulation

THP-1 cells were maintained in RPMI-1640 (P04-18047Pan Biotech,Germany) supplemented with 10% Fetal Bovine Serum (Gibco) in 5% CO_2_ in humidified environment at 37°C. For all the experiments, THP-1 monocytes were differentiated into macrophages with 50ng/ml PMA (Phorbol 12-myristate 13-acetate) (Sigma) for 16 hours followed by 24 hours of incubation in complete medium (Mehto et al., 2015; Sharma et al., 2016). The optimum concentration of protein for maximum activity was standardized using titration assay for MprA and MprA* (Supplementary Fig 2).The differentiated THP-1 cells were treated with 15 μg/ml of MprA and MprA* in all the subsequent experiments unless otherwise mentioned.

### Uptake and Intra-cellular detection of MprA and MprA*

MprA and MprA* were biotinylated with NHS-Biotin and subsequently conjugated with streptavidin-phycoerythrin (Strep-PE) (Chadha et al., 2015; Mehto et al., 2015; Bandyopadhyay et al., 2017). PMA-treated THP-1 cells were incubated with 15 μg/ml of biotin-streptavidin-phycoerythrin (PE) conjugated MprA and MprA*followed by dialysis to get rid of unbound biotin-streptavidin-phycoerythrin (PE) conjugate. Unstimulated THP-1 cells without any protein addition were taken as control. Uptake and internalization were scored at 1h interval at 60X on a Nikon C2 confocal microscope. Z-stack images were collected at 1μm intervals to score for internalization of the protein.

### Western blotting

Nuclear and cytoplasmic extracts were prepared from differentiated THP-1 cells using Nuclei EZ Prep kit (Sigma-NUC101) both from un-stimulated and MprA stimulated cells. 25μg of total protein was fractionated on SDS Polyacrylamide gel electrophoresis (PAGE). Lamin A/C antibody (Abcam) was used as nuclear marker and GAPDH was used as marker for whole cell extract. MprA was detected using Anti-His antibody (Invitrogen) and anti-mouse HRP-labelled secondary antibody (Santa Cruz). Following washing, the blots were developed using Luminol reagent (Santa Cruz). The intensity of the protein bands was measured and normalized to that of GAPDH.

### Estimation of secreted cytokine by ELISA

The differentiated THP-1 cells were checked for the levels of secreted cytokines in the culture supernatants by ELISA post-stimulation with MprA and MprA* for 24h.Unstimulated cells were taken as control. The levels of IFN-γ, TGF-β, IL-10, IL-17A and IL-6 were measured using the standard protocol mentioned in the e-Bioscience kit (San Diego, CA, USA).

### RNA extraction and expression analysis

RNA was isolated from 10^6^ cells following stimulation with MprA and MprA* for 24h using TRIzol (Invitrogen) method. Unstimulated cells were taken as control. 1μg of RNA was used for the cDNA preparation, using First strand cDNA synthesis kit (Fermentas K1612) qPCR was performed using Fast Start Universal Master mix (ROX) (Sigma) using exon spanning primers:-IL17A_Fp TGGAATCTCCACCGCAATGA, IL17A_Rp GCTGGATGGGGACAGAGTTC; TGFβ_Fp GATGTCACCGGAGTTGTGCG, TGFβ_Fp GTGAACCCGTTGATGTCCACTT; GAPDH_Fp CCAGGCGGCCAATACGAC, GAPDH_Rp AGCCTCCCGCTTCGCTCT. The values for specific genes were normalized to human *GAPDH* gene and fold change was calculated by ΔΔCt method.

### Apoptosis assay

THP-1 cells were stained with Annexin V-APC according to the manufacturer’s protocol(eBiosciences, USA) following stimulation with MprA and MprA*for 24h as described by Sharma et.al (2016)(Sharma et al., 2016).Unstimulated cells were taken as control. Approximately 10^6^cells were used in each group. Data acquisition and analysis was carried out on FACS Calibur (BD Biosciences).

### Phagosome-lysosome fusion

PMA differentiated THP-1 macrophages were stimulated with 15μg/ml MprA and MprA* for 24 hours, infected with GFP-*M.bovis* BCG at Multiplicity of infection (MOI) of 10, for 4 hours with or without pre-treatment with 2 μM of ionomycin for 45 min. The colocalization of GFP-*M.bovis* BCG with the lysosome was detected using lysotracker-Red (Invitrogen). Z stacking confocal imaging was performed with a Nikon C2 laser scan confocal microscope with a numerical aperture of 1.4, and a refractive index of 1.5 and 60X objective. The data were analysed using the Image J software (NIH USA) using Manders’ correlation coefficient method.

### Monitoring intracellular survival of *Mycobacterium*

Differentiated THP-1 cells were stimulated with MprA and MprA*. Unstimulated cells were taken as control. After 1h, cells were infected with 10 MOI of GFP labeled *M.bovis* BCG for 72h. Post-infection, the cells were thoroughly washed with 1X PBS to remove extracellular bacteria. The samples were mounted on slides with fluoroshield mounting medium containing DAPI. Single plane confocal images were acquired using a Nikon C2 laser scan confocal microscope. Data were analyzed using the NIS Elements Advanced Research software.Mean Fluorescence Intensities of 5 different fields from 3 independent experiments were measured.

### Statistical analysis

All statistical analyses were performed with Sigma plot 14 software, and the data were expressed as mean values with their standard deviations from 3 or more independent experiments. The results were analysed by non-parametric analysis of variance (ANOVA), followed by Tukey’s post hoc analysis. A p<0.05 was considered significant.

### Structural analysis of MprA and MprA*

In the absence of crystal structure for MprA, the effect of SNV in MprA protein was deciphered by generating the structure through homology modeling using I-TASSSER (Roy et al., 2010). The stereo-chemical properties of structure were analysed at SAVES (The Structure Analysis and Verification Server) and molecular dynamics (MD) simulation was used to optimize the atomic coordinates of MprA. The energy minimized structure was used to create mutation G70S (MprA*) with SPDBV (Johansson et al., 2012). To examine the effect of mutation (G70S) on the conformational dynamics of MprA, all atoms MD simulation was performed using the bio simulation package GROMACS 4.6.5 (Pronk et al., 2013) with the GROMOS 96 43A1 force field and SPC216 as water model.

The protein was placed centrally in the simulation box having dimension of 74.5 Å^3^ as x, y and z plane and the water molecules (~12291) padding around the protein. To neutralize the system, 0.15M Na+ and Cl□were added distal to protein in solvent and other parameters were defined as given earlier (Kumari, Idrees, et al., 2016; Kumari, Mishra, et al., 2016; Mishra et al., 2018). The prepared system was energy minimized using steepest descent and conjugant gradient, each run for 10000 steps.

Two ensemble processes NVT and NPT were used to equilibrate the system and PBC (periodic boundary conditions) was defined to x, y and z directions. During the simulation, temperature and pressure were maintained using Berendsen thermostat and Parrinello-Rahman pressure coupling, respectively. LINC algorithm (Hess et al., 1997) was used to constrain the bond lengths and hydrogen atoms. The electrostatic interactions were evaluated by PME (particle mesh Ewald) method (Darden et al., 1993) and LJ potential was used for van der Waals interactions. Using the NPT ensemble, two independent runs were performed on MprA and MprA* for the period of 250 nanosecond (ns) at 300 Kelvin (K). All the coordinates, trajectory, energy and velocity were saved at the time interval of 5 picosecond (ps). The obtained trajectories were analysed using the GROMACS utilities. XMGrace was used for the analytical plots.

## Results

We identified a SNV (Single Nucleotide Variation) in genomic position G_1097023_A in *mprA* gene that led to the substitution of an amino acid, glycine to serine at 70^th^ position (G70S) in one of our clinical isolates VPCI591. This variation was detected in 3% of the global clinical isolates, as analyzed from GMTV database (Chernyaeva et al., 2014) and tbVar database(Joshi et al., 2014) containing about 1553 global clinical isolates of *M.tuberculosis.* In the light of the identification of MprA as late antigen and its role implied in persistence of *M. tuberculosis,* we examined the effect SNP.

### Nuclear localization of MprA

In order to analyse the effect of the SNV, the uptake and intracellular localization of the protein in THP-1 macrophages were examined. MprA and MprA* tagged with biotin-sterptavidin-phycoerythrin (PE) were internalized and completely localized in the nucleus of the THP-1 macrophages within 1 hour of incubation, as assessed by confocal microscopy (Figure 1A). THP-1 cells without the addition of any protein were taken as the control where only DAPI stained nuclei were seen.

**Figure 1:**
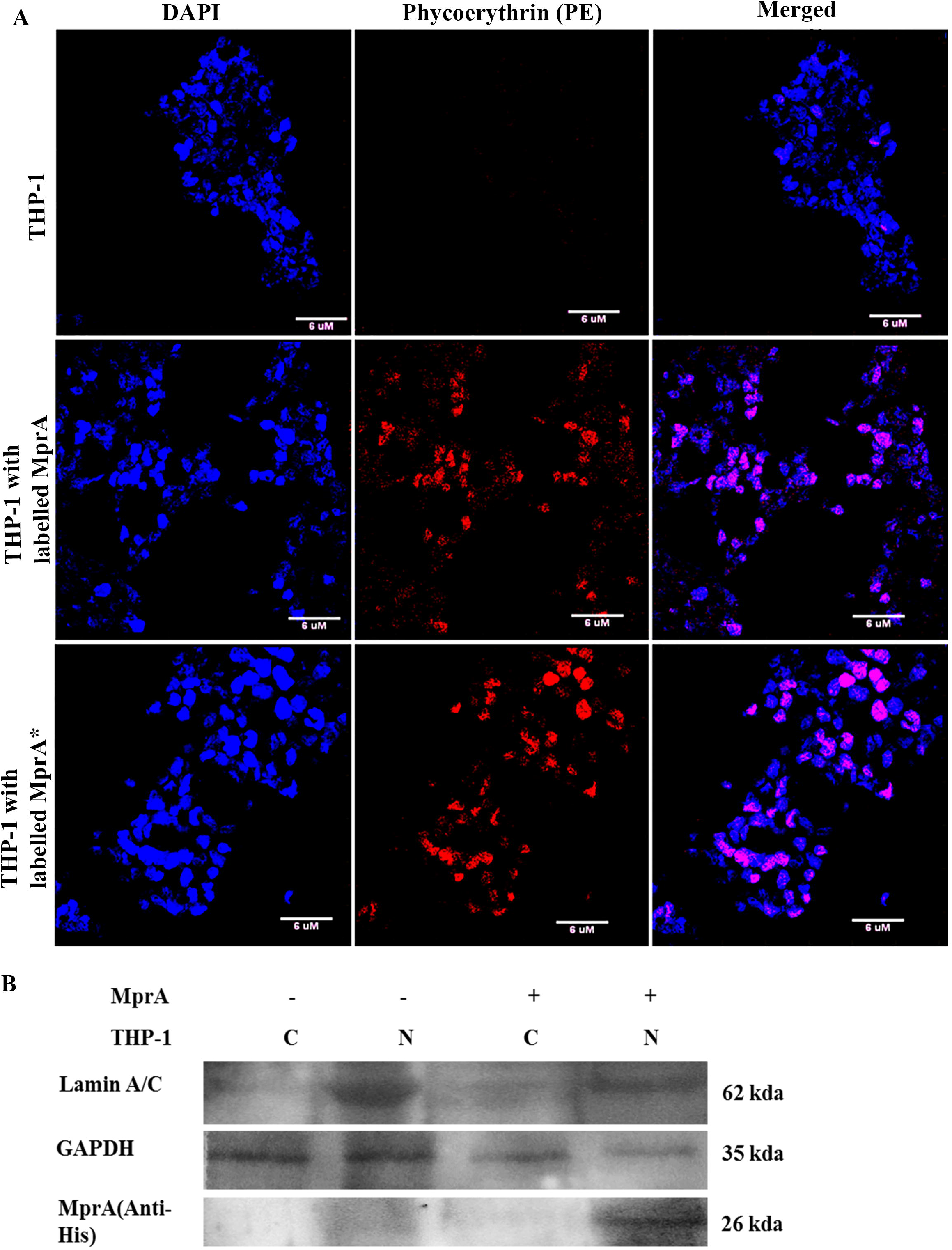
MprA and MprA* localise in the nucleus of THP-1 macrophages. THP-1 macrophages were incubated with 15 μg MprA and MprA* for 1 h. Localization was monitored by confocal microscopy (Panel A) or Western blotting (Panel B) as detailed in Materials and methods. In Panel A top row shows differentiated THP-1 cells used as control without the addition of MprA or MprA*, middle row shows THP-1 cells stimulated with MprA-Biotin-Strep-PE, bottom row shows THP-1 cells stimulated with MprA*-Biotin-Strep-PE. Merged images show MprA/MprA*-Biotin-Strep-PE localization in the nuclei. One of three experiments is shown. For Panel B, Nuclear (N) and cytoplasmic(C) fractions were analysed for the presence of MprA. Lamin was used as marker for nuclear extract and GAPDH used as marker for whole cell extract. The antibody used is indicated on the left. Nuclear and cytoplasmic extract from THP-1 cells with no MprA addition was used control.

The cytoplasm and nuclear fractions of THP-1 cells stimulated by MprA was analysed by Western blot using anti-His antibody. The MprA protein was detected in the nuclear extract and not in the cytoplasm (Figure 1B). Similar results were obtained with MprA*(data not shown). Lamin A/C was used as the nuclear marker and GAPDH was used as marker for whole cell extract (Figure 1B).

### MprA and MprA* modulate defence responses of macrophages

We examined the role of MprA and MprA* in influencing the defence responses of THP-1 macrophages in terms of cytokine production, apoptosis, phagosome-lysosome fusion and survival of *M.tuberculosis* inside macrophages.

Earlier studies have shown several *M. tuberculosis* proteins, including MprA, to affect multiple aspects of host immune response (Zahrt & Deretic, 2001; Gupta et al., 2010; Pang et al., 2013). Therefore, we analysed the effect of MprA and MprA* on the levels of different cytokines in THP-1 cells by ELISA. Among the several cytokines checked for their level, the level of IL-17A,TGF-β and IL-10 was found to be elevated (Figure 2, Panel A-C).Further, the variant MprA*, brings about significantly higher alteration in IL-17A and TGF-β level compared to MprA. No significant change in levels of IFN-γ, and IL-6 levels was observed (data not shown). The increase in IL-17A and TGF-β is concurrent with the increase in the transcription of the respective genes (Figure 2, Panel D,E). This in turn correlates with the internalization of MprA/MprA* that needs further substantiation.

**Figure 2:**
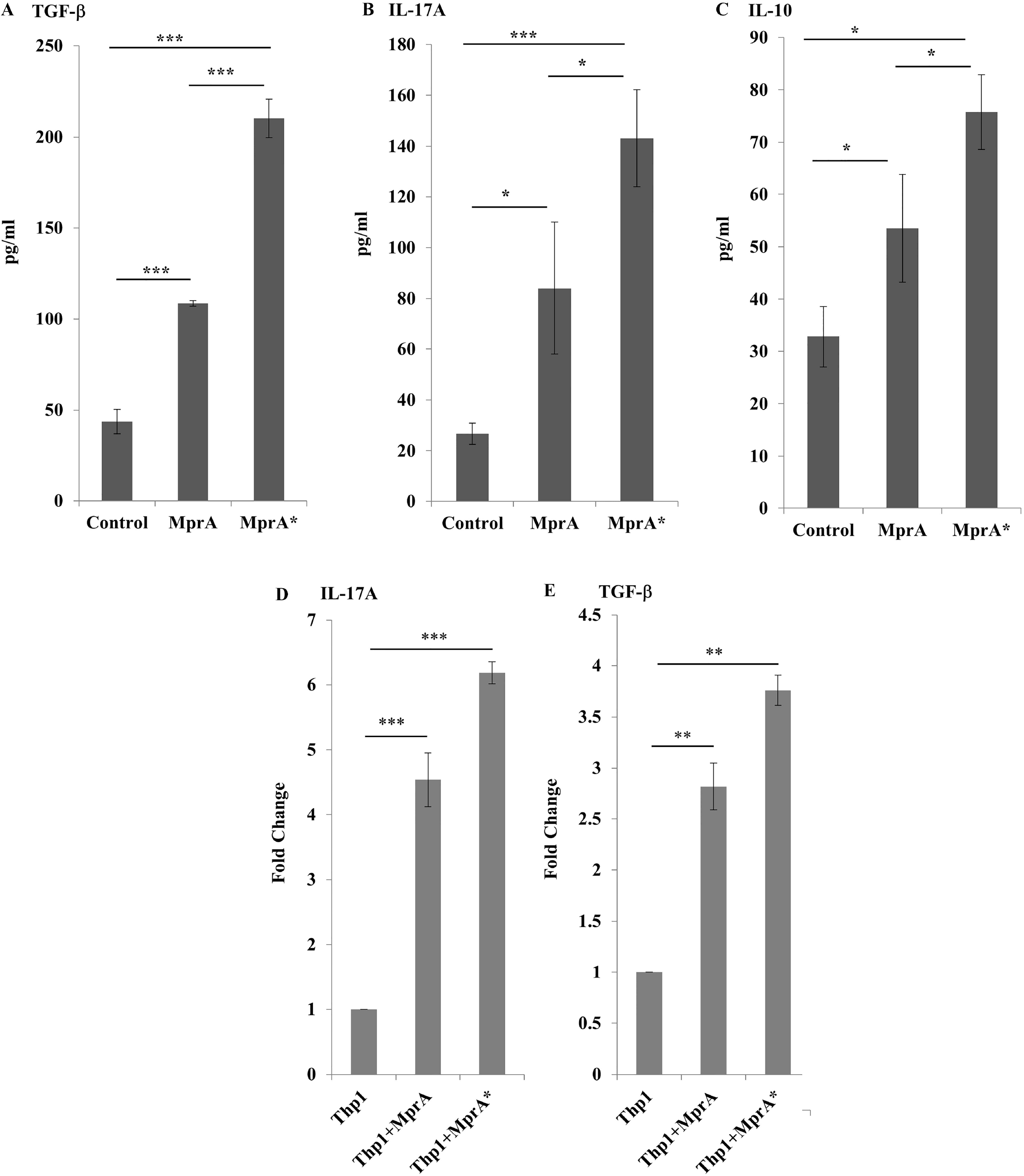
MprA and MprA* modulate cytokine profiles in macrophages. THP-1 macrophages were incubated with 15 μg of MprA and MprA* for 24h. The level of cytokines was measured in the culture supernatants by ELISA (Panel A-C) or q-PCR (Panel D-E). Data represent means ± SD of 3 independent experiments. ANOVA was performed with 95% CI. For groups Control, MprA and MprA* the results were analysed by one way ANOVA followed by Tukey’s post hoc multiple comparison test, *(*P* ≤ 0.05), ***(P ≤ 0.001). MprA* shows higher increase of cytokines than MprA. For Panels D-E, Also total RNA was isolated using TRIzol method from stimulated with MprA and MprA* for 24h. The transcript levels of IL-17A and TGF-β measured by q-PCR is shown in terms of fold change. Data from three independent experiments are shown. For groups Control, MprA and MprA* the results were analysed by one way ANOVA followed by Tukey’s post hoc multiple comparison test, **(P ≤ 0.01), ***(P ≤ 0.001).

### MprA promotes macrophage survival

A key strategy used by *M. tuberculosis* to establish long-term infection in the macrophage is to prolong its survival by inhibiting apoptosis (Mehto et al., 2015). As shown in Figure 3, stimulation of macrophages with MprA and more significantly MprA*, decreased Annexin V levels thus contributing to the increased survival of the cells that may relate to the long term persistent infection.

**Figure 3:**
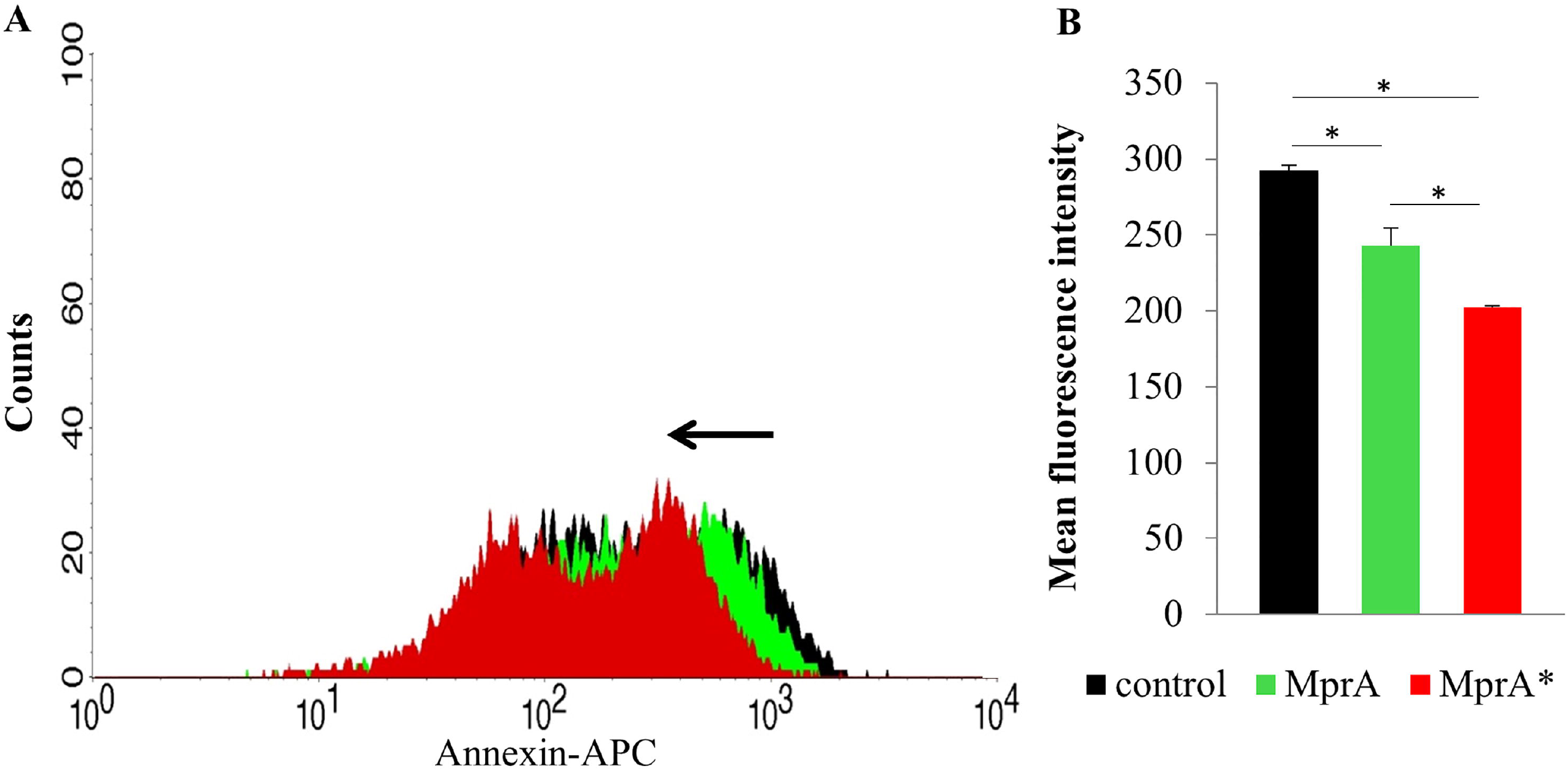
MprA and MprA* modulates apoptosis. THP-1 macrophages were incubated with 15 μg of MprA and MprA* for 24h and stained with Annexin V-APC. The Black histogram shows unstimulated cells, green histogram shown cells stimulated with MprA and Red histogram shown cells stimulated with MprA*. Panel A shows the distribution and a backward shift is observed, Bar graphs in Panel B shows mean fluorescent intensity of the indicated groups represented as Mean ± S.D. Results of three independent experiments are shown for the groups control (unstimulated cells), MprA and MprA*.The results were analysed by one way ANOVA followed by Tukey’s post hoc multiple comparison test, *(P ≤ 0.05).

The fusion of phagosomes with lysosomes leads to clearing of the pathogen. To avoid the distal immune responses and to promote long-term persistence of infection, the fusion of phagosomes and lysosomes is prevented. We investigated phago-lysosome fusion in presence of MprA/MprA* using *M.bovis* BCG.

As *Mycobacteria* inhibit phagosome-lysosome fusion, we first created conditions that would promote phago-lysosome fusion. It is well established that increased calcium concentration in infected cells promote this fusion and *M.tuberculosis* down modulates calcium levels upon infection(Malik et al., 2000; Malik et al., 2003). We recently showed that *M.tuberculosis* protein Rv3529c inhibits the fusion of phagosomes with lysosomes that is promoted by increased influx of calcium mediated by the calcium ionophore, ionomycin (Bandyopadhyay et al., 2017). We therefore, used the same strategy to investigate the role of MprA and MprA* in modulating phago-lysosome fusion.

Infection of *M.bovis* BCG did not show appreciable phago-lysosome fusion, however treatment with ionomycin induced significant levels of phago-lysosome fusion. Under these conditions, incubation of macrophages with either MprA or MprA* significantly inhibited ionomycin induced fusion of *M.bovis* BCG phagosomes with lysosomes which is also evident from Manders’ correlation coefficient calculated for the samples (Figure 4A,B). This suggests a suppressive role for MprA and MprA* in reducing phago-lysosome fusion to enhance the survival of the pathogen inside the THP-1 macrophages. MprA* reduced the fusion frequency more than MprA (Figure 4A,B).

**Figure 4:**
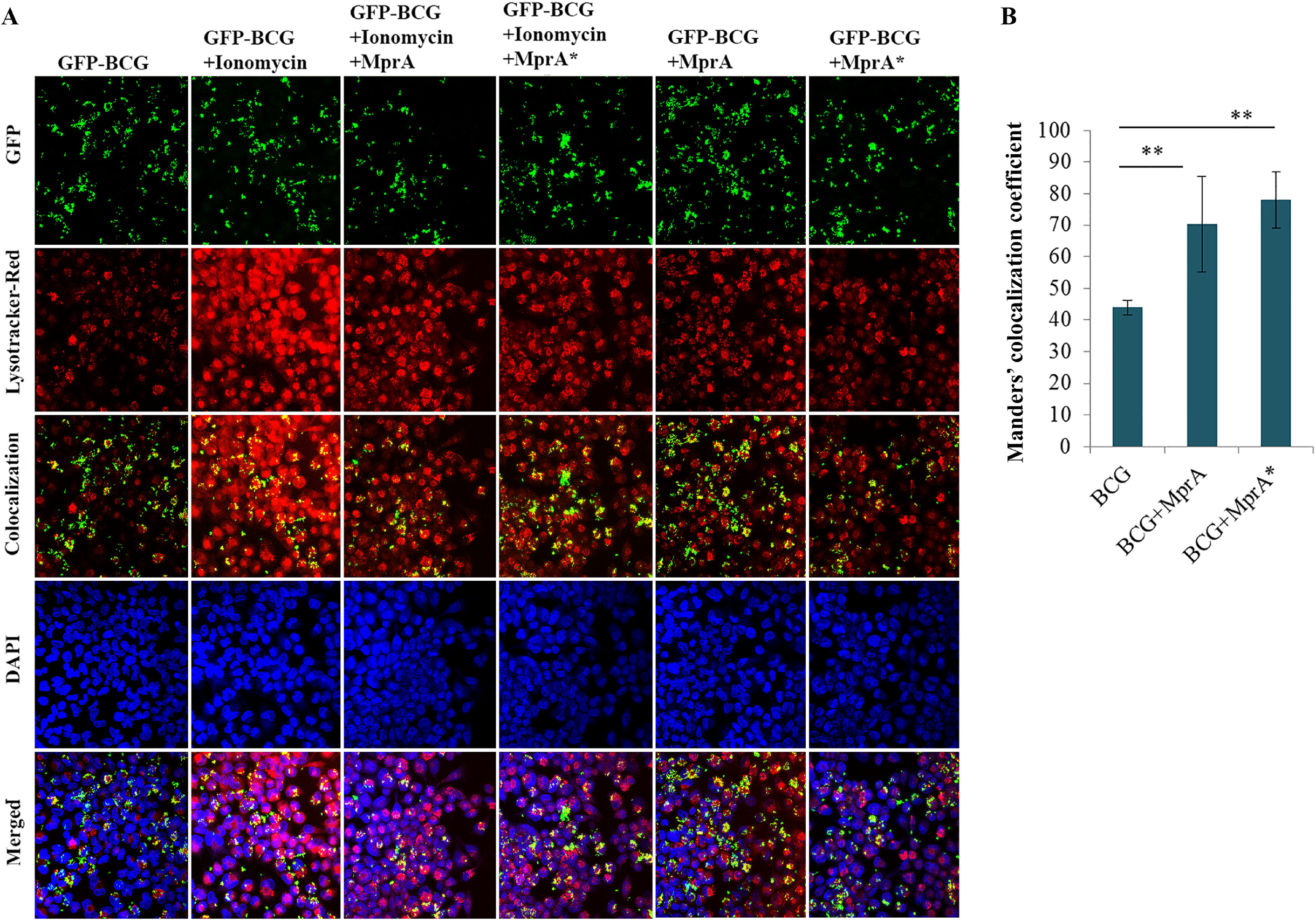
MprA and MprA* inhibit Phago-lysosome fusion in macrophages. THP-1 macrophages were incubated with 15 μg of MprA and MprA* for 24h followed by infection with 10 MOI GFP-BCG for 4h. For some groups cells were also treated with 2μM ionomycin (see Materials and Methods). Cells were stained using Lysotracker Red to stain lysosomes. Confocal microscopy was performed using Nikon C2 laser confocal microscope. Green represents GFP-BCG, Red represents Lysosomes, Blue represents Nucleus stained with DAPI. Scale bar in the lower column applies to all of the images. B. Bar graphs represent Manders’ correlation coefficient for colocalization of GFP-BCG with lysosomes from the Z stack.

The downstream effect of apoptosis inhibition and reduced phago-lysosome fusion in infected cells could increase bacterial survival. We therefore, investigated if MprA and MprA* would influence bacterial survival inside the macrophages. To that end, following stimulation and incubation of the macrophages with MprA and MprA*, the cells were infected with GFP tagged *M. bovis* BCG for 48h. On comparison with the control, the macrophages stimulated with MprA and MprA* supported enhanced survival of the bacteria (Figure 5). The mean percentage localization was measured as an indicator of the bacterial load inside the cells.

**Figure 5:**
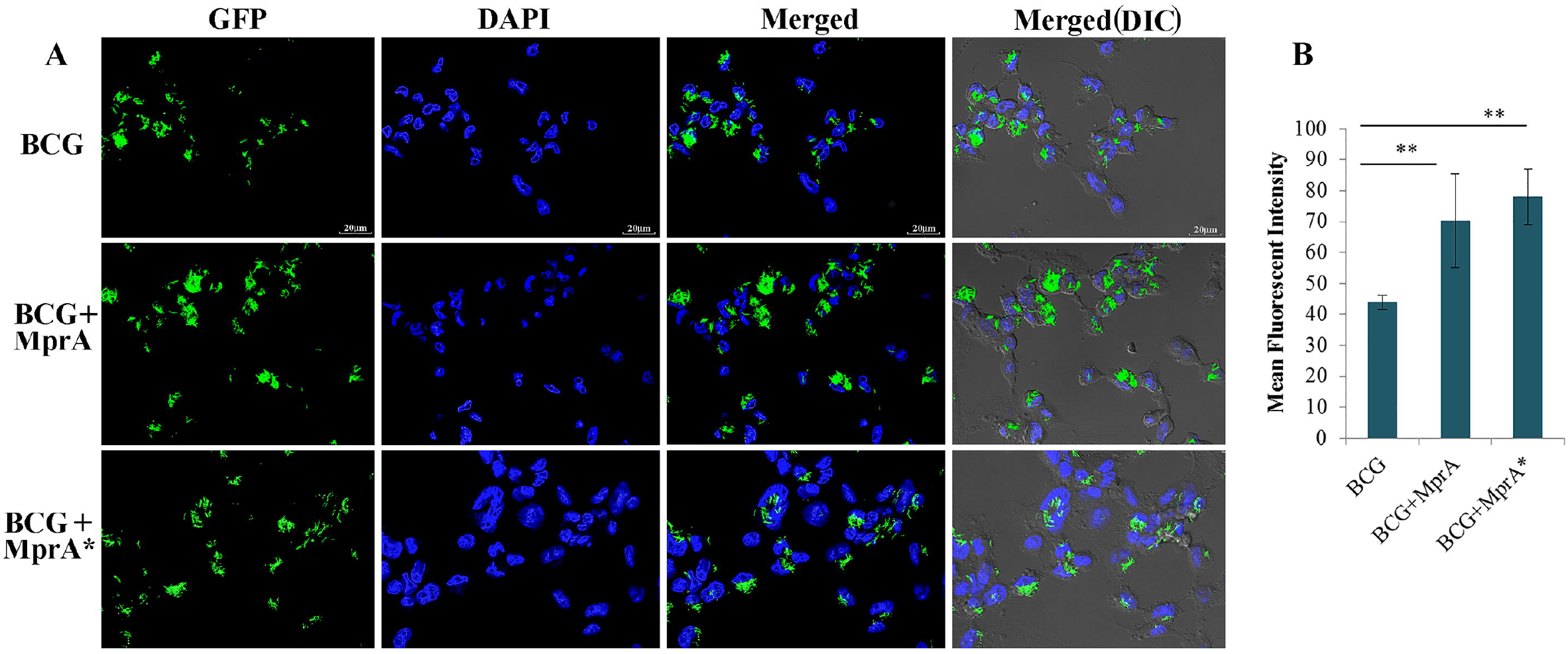
MprA and MprA* aid the survival of pathogen inside macrophages. THP-1 macrophages were incubated with 15 μg of MprA and MprA* for 24h followed by infection with 10 MOI GFP-BCG for 72h. Cells were fixed and processed for confocal microscopy. Green represents GFP-BCG while Blue represents nucleus stained with DAPI. Bar graphs in the figure show the Mean Fluorescence Intensities of 5 different fields from 3 independent experiments.Scale bar in the upper column applies to all of the images B. The bar graph indicates mean fluorescent intensity of groups in cells infected with BCG, BCG+ MprA and BCG+ MprA*.The results were analysed by one way ANOVA followed by Tukey’s post hoc multiple comparison test**(P ≤ 0.01).

### Implication of non-synonymous substitution on the structure of MprA

To determine the effect of the SNV on the conformational dynamics and stability of MprA, we examined the time evolution plot of C^α^-RMSD as shown in Figure 6A. We observed that RMSD trajectory of MprA attains the equilibrium ~100 ns and remains stable till the simulation is completed at 250 ns. In comparison to MprA, RMSD plot of MprA* shows several consecutive drifts in trajectory during 0-150 ns, indicating the reduction in stability of the structure of the variant protein. However, we can see the drop-down in drifts around ~150-250 ns during the last 100 ns of simulation, which indicates the stable conformational dynamics of MprA*.

**Figure 6:**
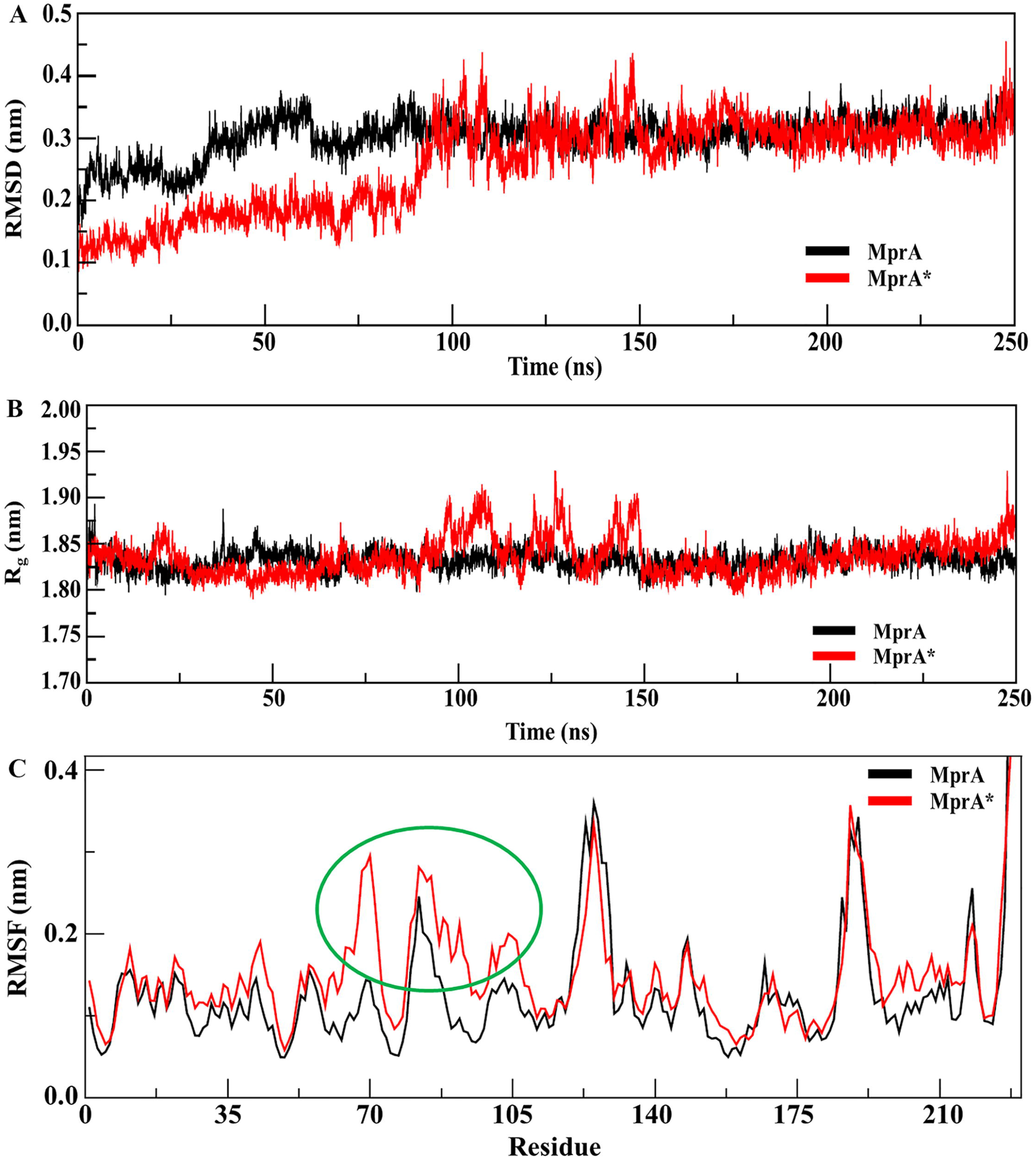
Molecular Dynamic (MD) simulation for structural comparison between MprA and MprA*. A. Backbone RMSD plots of MprA (black) and MprA* (red) in water at 300 K during 250 nano second MD simulation are shown. B.Radius of gyration (Rg) MprA (black) and MprA*(red) during 250 nano second of MD simulation. C.Time-average RMSF plot per residue of MprA (black) and MprA*(red) during 250 nano second of MD simulation are shown. Fluctuations observed for MprA* at (G70S) and adjacent residues are shown with green circle.

We also examined the structural stability of MprA and MprA* using the conformational order parameter R_g_ which defines the structural integrity and the compactness of a protein. The time evolution plot of R_g_ for all Cα-atoms is shown in Figure 6B. The structure of MprA remains stable around R_g_ value 1.83±0.01 nm, whereas, several drifts in R_g_ trajectory of MprA* can be seen during ~75-175 ns. The higher fluctuation in structure with the drifts of R_g_ ≥ 0.1 nm at 100-150 ns provide a clear evidence of less stable structure of MprA* as compared to MprA. The time average change in fluctuation of each residue within the RMSF plot was analysed to examine the effect of MprA* on the dynamics of the local structure of the protein (Figure 6C).

To understand the effect on flexible regions of wild type MprA, the average position fluctuation of amino acid residues in a dynamic system was analysed. It indicates that the amino acid alteration in MprA*, contributes to the fluctuation in the secondary and tertiary structure. The result showed that the alteration leads to increased fluctuation of Cα backbone residues at the 70^th^ position and further to the amino acids around G70S in MprA*. This investigation suggested that the protein MprA* shows significant alteration of about 0.2nm and specifies the consistency with RMSD and Rg data. However no major structural change was observed.

## Discussion

The detection of single nucleotide variation(SNV) in clinical isolates is reported in a large number of cases (Sandgren et al., 2009), however, the functional implication of SNVs is known in only in limited number of cases mainly in genes conferring drug resistance. In the present study we focused on the functional outcomes of MprA (*Rv0981*) and MprA* (G70S) present in the multidrug resistant clinical isolate VPCI 591. The importance of MprA in the survival and persistence is well known. As a member of the OmpR group of transcription factor, it is known to have DNA binding activity and the binding domain of MprA in *M.tuberculosis* has also been identified(Pang et al., 2013; Banerjee et al., 2016). However, the SNV in MprA* does not map in this region.

The localization of MprA in the macrophage nucleus is a unique feature. However, there are examples where mycobacterial proteins are transported to the eukaryotic nucleus by other proteins. For instance, PPE2(*Rv0256c*) which has a leucine zipper DNA-binding motif, is translocated into the macrophage nucleus through the classical importin α/β pathway (Bhat et al., 2017). Rv3423.1, a mycobacterial histone acetyltransferase, is known to co-localize with chromatin in macrophage nucleus while it was secreted into the culture medium by the virulent strain of the bacteria (Jose et al., 2016).The authors demonstrated that Rv3423.1 protein localizes with the chromatin and brings about histone acetylation. In this case also no nuclear localization signal was detected and it was speculated that it utilizes host protein to hitchhike into the nucleus. Similarly, (Sharma et al., 2015) reported the localization of Rv2966c protein with host chromatin to bring about non-CpG methylation. In this case, the N-terminal of the protein was seen to be important for nuclear localization. However we did not find any nuclear localization signal in the protein MprA.

The antigenic property of MprA and also the detection of RNA following *M.tuberculosis* H37Rv in mice are reported by (Gupta et al., 2010). MprA affects persistent infection through the regulation of several genes in *M.tuberculosis* (Zahrt et al., 2003; He et al., 2006; Pang et al., 2007). It is up-regulated following infection in human macrophage cells (Haydel & Clark-Curtiss, 2004). MprA and other antigens of *M.tuberculosis* lead to varied profiles of the cytokines, overall contributing to the immune-suppression observed on *M.tuberculosis* infection(Gupta et al., 2010). The nuclear localization of MprA/MprA* observed in our study strongly suggests that proteins of the pathogen may modulate the expression of host genes. The transcriptional up regulation of IL-17A, TGF-β and IL-10, raises the possibility of involvement of MprA/MprA* in host gene regulation, either directly or indirectly. However we failed to detect any canonical or variant DNA motif for binding of MprA in the upstream regions of IL-17A, TGF-β and IL-10.

The enhanced level of IL-17A, TGF-β and IL-10 indicates a propensity of MprA and MprA* to drive a TH-17 response as well as push the response towards a suppressive phenotype with higher levels of TGF-β and IL-10. TGF-β is required for both TH-17 responses as well as promoting regulatory T cell responses. Recent reports have indicated a dual role for IL-17 in mediating both protective as well as suppressive responses towards *Streptococcus pneumoniaeinfection* (Ritchie et al., 2018) and also immune protective responses in hyper-virulent tuberculosis infection (Gopal et al., 2014). IL-17 is seen as a mediator in the pathology in auto-immune diseases as well. It is also known to maintain granuloma integrity where it contributes to a protective response against tuberculosis infection (Umemura et al., 2007). It would be interesting to examine if IL-17 plays a similar role in mycobacterial infections. The fact that no significant change was observed for IFN-γ, and IL-6, indicates that MprA and MprA* may not regulate the responses mediated by these cytokines. Gupta et al.,(Gupta et al., 2010) have shown compartmentalization of functions of several *M.tuberculosis* proteins expressed inside the macrophages, wherein the responses mediated by one protein are not duplicated by the other. Instead, a coordinated function of all the proteins in a time dependent manner ensures that suppressive responses are promoted and maintained at sites of *M.tuberculosis* infection. IFN-γ is a pro-inflammatory cytokine and since neither MprA nor MprA* induced its production clearly point towards attenuating protective responses. While IL-10 is a suppressive cytokine, the route of suppression mediated by MprA/ MprA* may be direct or via TGFβ. Further analyses of T cell responses elicited by MprA or MprA* would give additional insights into their role in immune suppression.

The fusion of phagosomes with lysosome is an important step in the host protection, as it leads to the presentation of the antigens from the pathogen leading to effective immune response. The evasion of apoptosis and the failure of phago-lysosome fusion and further increase of bacterial load observed on internalization of MprA, converge to the conclusion that MprA plays a role in immune suppressive responses and adds support to the findings of Gupta et al., (2010). A similar phenomenon was seen to be more pronounced in case of MprA*. Both show reduced fusion of bacteria with lysosomes in the presence of ionomycin (Ca^2+^ inducer of phago-lysosome fusion) which has been depicted in the form of Manders’ overlap coefficient (MOC) of colocalization index. It has also been demonstrated recently that voltage-gated calcium channel regulated calcium influx and homeostasis has an important role in regulating phago-lysosome fusion and can prolong the survival of the pathogen in the macrophage (Malik et al., 2000; Sharma et al., 2016). It is known that MprA has a role in persistence of infection (Zahrt & Deretic, 2001). Our result suggests that MprA and MprA* show apoptosis-rescue mechanism on THP-1 differentiated macrophages correlating with the immune suppressive behaviour of the protein.

In the context of the above experiments, we examined if there is any major structural alteration of MprA* attributable to the SNV. From the RMSF (Root mean square fluctuation) analysis MprA* has shown fluctuation at 70th position of the protein and the nearby residues. However, no major structural alteration was seen in MprA* with respect to the wild type MprA.

We conclude that, the current study has provided evidence of the nuclear localization of the pathogen protein that may lead to modulation of transcription of host genes, which can result in diverting the host machinery in favour of the pathogen. In addition, our results show that single nucleotide polymorphisms are not always benign, but SNVs can affect the function of mycobacterial proteins at a quantitative level.

## Acknowledgements

KN and VB acknowledge UGC Special Assistance Program (UGC-SAP-II) and DU-DST PURSE Grant. We acknowledge DBT Bioinformatics facility at ACBR. The authors acknowledge Dr.Richa Arya (ACBR) and Dr.Akanksha Verma (ACBR), for advice and help with the confocal experiments. KB acknowledges CSIR for Senior Research fellowship.

---

**Supplementary Fig 1: Coomassie staining of purified MprA and MprA*.** SDS-PAGE showing purified protein MprA(A) and MprA*(B). Lane 1-protein marker, Lane 2-Uninduced, Lane 3-wash 1, Lane 4-wash 2, Lane 5,6,7,8-250mM Imidazole successive eluting fraction, Lane 9-soluble fraction, Lane 10-Ni-NTA slurry.

**Supplementary Fig 2: Standardisation of the optimum concentration of MprA and MprA* for stimulation.** MprA and MprA*concentration was optimized using titration assay for IL-10 and TGFβ and the results are represented for MprA (A,B) and MprA* (C,D) as bar diagram.

