## Supplementary figures and images for "Modulation of macrophage defense responses by Mycobacterial persistence protein MprA (*Rv0981*) in human THP-1 cells: effect of single amino acid variation on host-pathogen interactions"

### Supplementery Fig 1

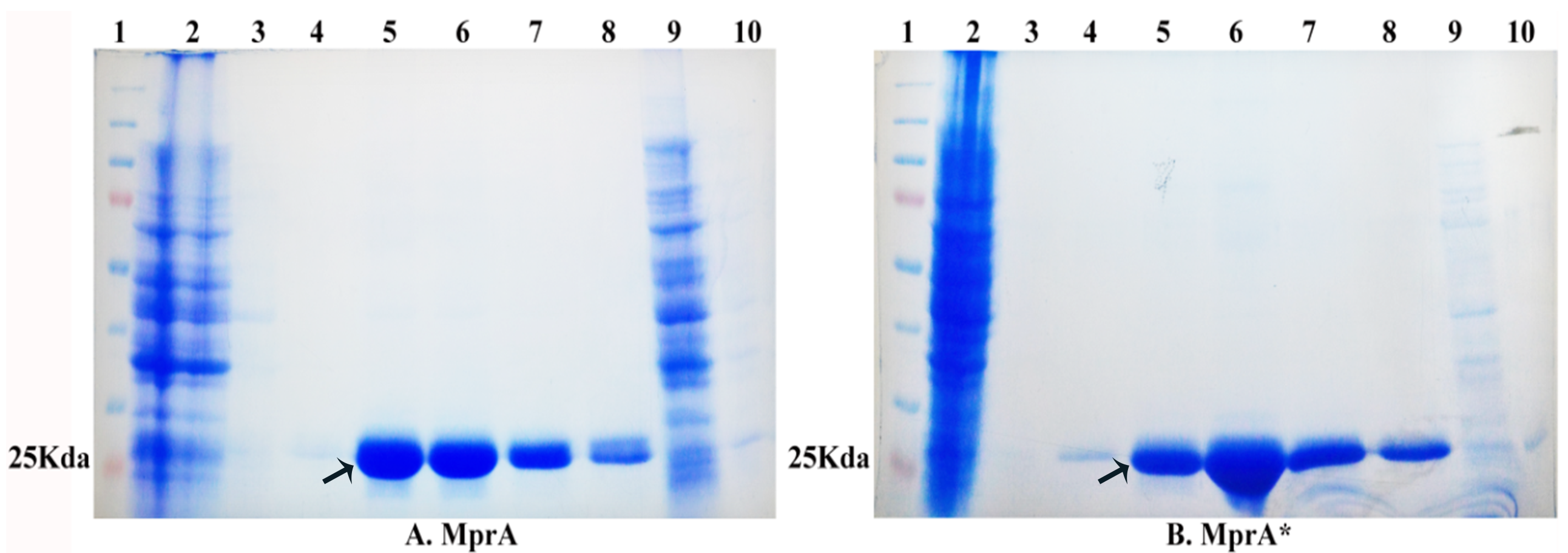

### Supplementery Fig 2

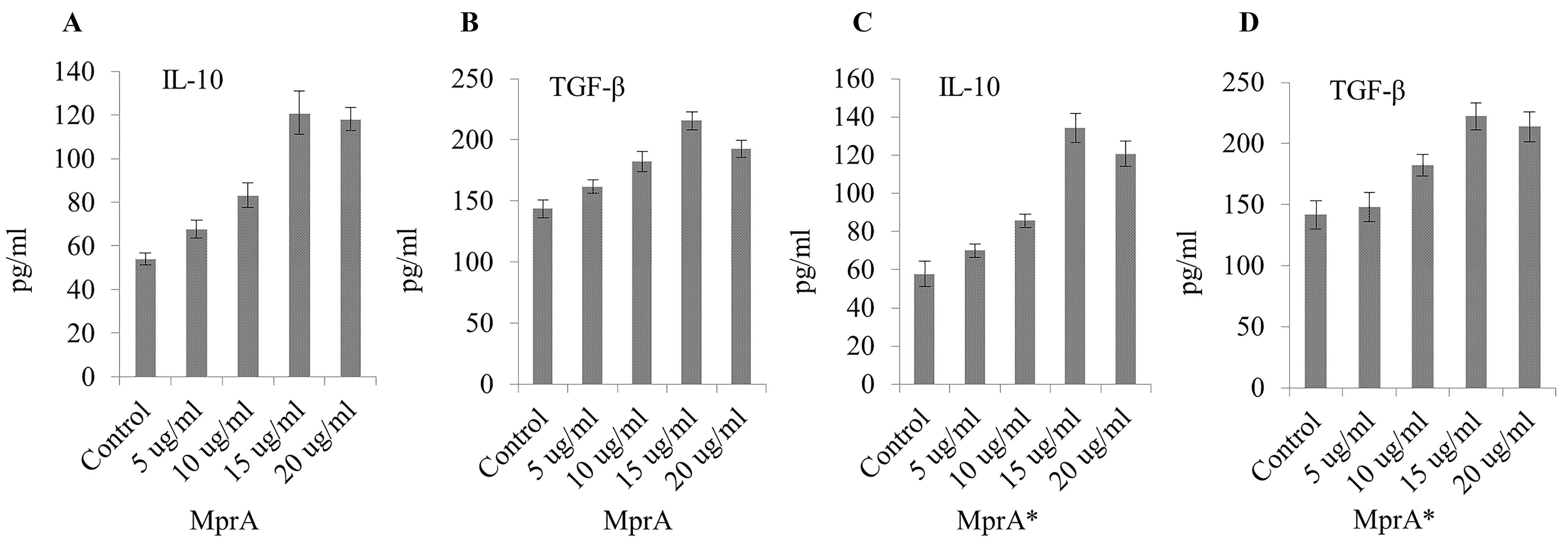
